# A genetic screen for regulators of muscle morphogenesis in *Drosophila*

**DOI:** 10.1101/2021.03.04.434006

**Authors:** Tiffany Ou, Gary Huang, Beth Wilson, Paul Gontarz, James B. Skeath, Aaron N. Johnson

## Abstract

The mechanisms that determine the final topology of skeletal muscles remain largely unknown. We have been developing *Drosophila* body wall musculature as a model to identify and characterize the pathways that control muscle size, shape, and orientation during embryogenesis (Johnson et al., 2013; Williams et al., 2015; Yang et al., 2020a; Yang et al., 2020b). Our working model argues muscle morphogenesis is regulated by (1) extracellular guidance cues that direct muscle cells toward muscle attachment sites, and (2) contact dependent interactions between muscles and tendons. While we have identified several pathways that regulate muscle morphogenesis, our understanding is far from complete. Here we report the results of a recent EMS-based forward genetic screen that identified a myriad of loci not previously associated with muscle morphogenesis. We recovered new alleles of known muscle morphogenesis genes, including *bsd, kon, ths*, and *tum*, arguing our screening strategy was effective and efficient. We also identified and sequenced new alleles of *salm*, *barr*, and *ptc* that presumably disrupt independent pathways directing muscle morphogenesis. Equally as important, our screen shows that at least 11 morphogenetic loci remain to be identified. This screen has developed exciting new tools to study muscle morphogenesis, which may provide future insights into the mechanisms that determine skeletal muscle topology.

## Introduction

The mechanisms that regulate skeletal muscle morphogenesis have been remarkably understudied across Metazoa. Body wall muscles in the *Drosophila* embryo form a stereotyped pattern with 30 distinct muscles arranged in a spectacular compilation of longitudinal, acute, and oblique orientations (Fig. 1). Remarkably, the final muscle pattern shows great diversity along the dorsal-ventral axis, but the pattern is invariant from segment to segment along the anterior-posterior axis (Bate, 1990). A single embryonic segment must therefore house the essential morphogenetic information required to orchestrate the unique arrangement and functional morphogenesis of 30 individual muscles.

**Figure 1.**
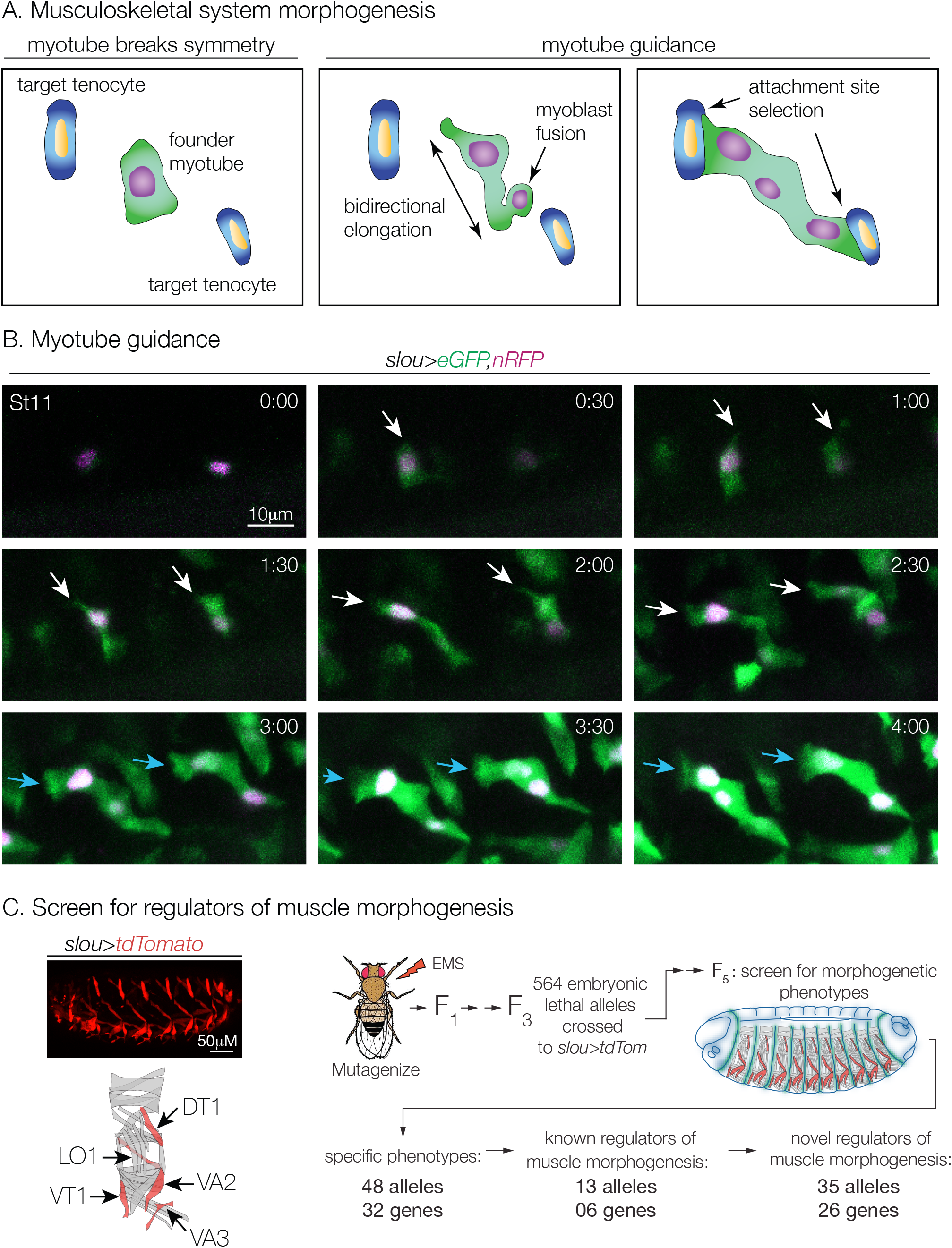
A genetic screen for regulators of muscle morphogenesis. (A) Diagram of *Drosophila* embryonic body wall muscle development. (Left) After cell fate specification, founder myoblasts break symmetry and become founder myotubes. (Right) Myotube leading edges elongate toward tenocytes (tendon precursors) that house the muscle attachment sites. Fusion competent myoblasts will fuse with founder myotubes during elongation. Each myotube leading edge ultimately selects the correct muscle attachment site. Myotube guidance encompasses both leading edge extension/navigation and attachment site selection. (B) Live imaging of myotube guidance. Two founder LO1 myotubes in neighboring segments expressed cytoplasmic eGFP (green) and nuclear RFP (violet) under the control of *slou.Gal4.* Arrows point to myotube leading edges during elongation (white) and during attachment site selection (blue). (#:##) hr:min. (C) Screening strategy. EMS-induced embryonic lethal alleles were identified and crossed with *slou>tdTomato* to visualize the morphology of the Longitudinal Oblique 1 (LO1), Dorsal Transverse 1 (DT1), Ventral Transverse 1 (VT1), Ventral Acute 2 (VA2), and Ventral Acute 3 (VA3) muscles in homozygous mutant embryos.

Mesodermal muscle precursors, known as myotubes, extend asymmetrical, amoeboid projections that navigate across the embryonic segment to contact pre-determined muscle attachment sites on ectodermal tendon precursors at the segment border. The mesenchymal myotube then establishes a myotendinous junction, regains symmetry, and acquires its final, functional orientation (Fig. 1A, B). Myotube guidance refers to the combined cellular processes of leading edge navigation and attachment site selection. During myotube guidance, morphogenetic information is transmitted to the muscle precursors through a bipartite system (Yang et al., 2020b). Short-range secreted signals from the ectoderm provide navigational cues, which direct ameboid projections toward specific muscle attachment sites. After the projections arrive at the segment border, a second type of morphogenetic information, which is presumed to reside in the tendon precursors, ensures the myotube selects the correct muscle attachment site. Incredibly, this bipartite information system ensures that individual cells from two distinct deterministic cell populations, which are specified in separate germ layers, locate one another with high fidelity during embryogenesis.

Our work to date suggested key molecules that provide or interpret morphogenetic information during myotube guidance remain to be discovered, so we screened for novel regulators of muscle morphogenesis. To extend previous muscle screens (Chen et al., 2008; Johnson et al., 2013), we used the shape and orientation of just 5 of the 30 muscles per segment to identify and classify muscle phenotypes at high resolution. By focusing on a subset of muscles, we successfully obtained new alleles in 26 genes that were not previously known to regulate embryonic muscle morphogenesis (Fig. 1C). We mapped 8 alleles to causative point mutations in 4 distinct loci that encode the zinc-finger transcription factor Spalt major (Salm), the chromatin binding protein Barren (Barr), the Hedgehog (Hh) signaling regulator Patched (Ptc), and the serine/threonine (ser/thr) kinase Back seat driver (Bsd). The reagents generated from this screen will provide novel inroads toward understanding the molecular mechanisms of muscle morphogenesis and myotube guidance.

## Materials and Methods

### Primary screen

To generate screening stocks, flies were isogenized on chromosome II and males were treated with 25mM EMS as described (Asburner et al., 2005). Mutagenized males were bred as shown:

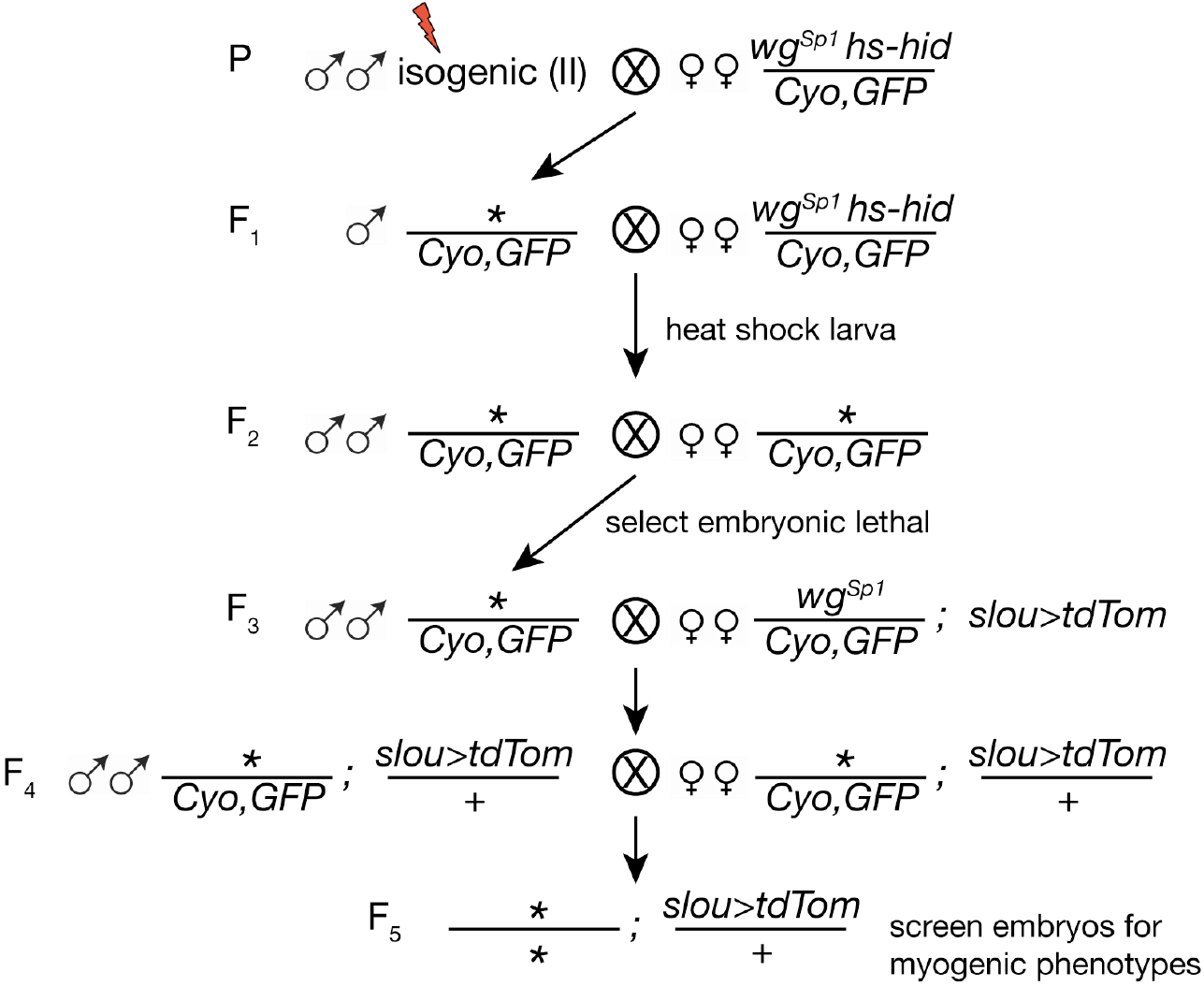

0-24hr F_5_ embryos were collected on grape plates, dechorionated, transferred to glass slides, and *slou>tdTom* expressing muscles were imaged in live embryos. *slou.Gal4* is active in 5 distinct muscles per segment (Fig. 1C). The presence and morphology of individual s*lou.Gal4* expressing muscles was used to assign mutations to phenotypic classes (see Table 1). At least 5 homozygous embryos were scored per line. 58 lines with myogenic phenotypes were recovered from the primary screen.

**Table 1.**
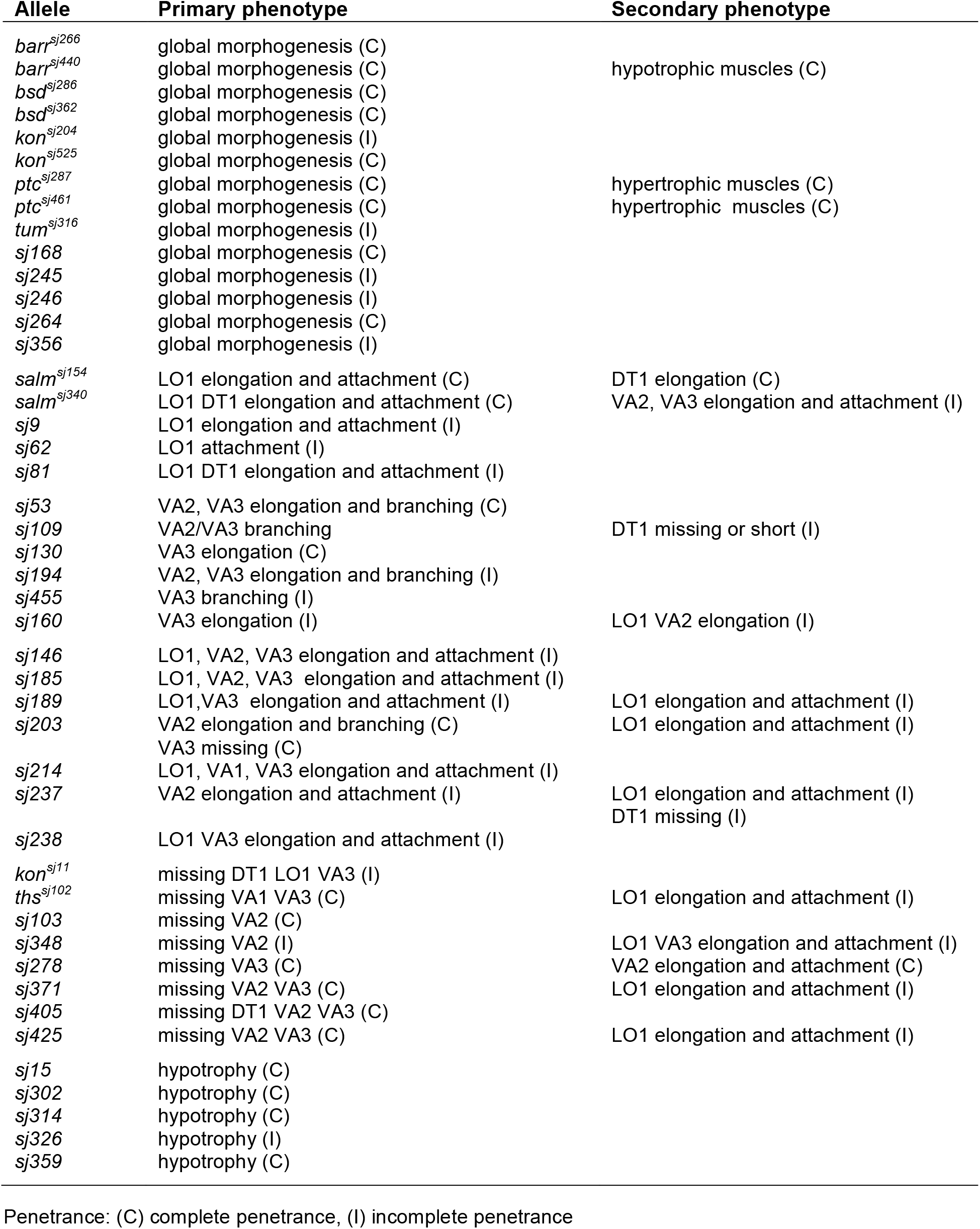
Phenotypic classes of myogenic alleles.

### Allele nomenclature

At the conclusion of the primary screen, each line was assigned a unique allele identifier of *sj###*. For mapped alleles, the unique identifier is superscripted with the gene symbol (e.g. *salm*^*sj154*^). Unmapped alleles are referenced by the unique allele identifier in figures and tables (e.g. *sj168*).

### Complementation analysis

The 58 lines with muscle phenotypes were inter-crossed, and transheterozygote adult viability was used for initial complementation analysis. Lines that failed to complement were further analyzed for transheterozygous embryonic phenotypes to confirm the non-complementing alleles disrupted myogenesis.

The 58 lines with muscle phenotypes were also crossed to lethal alleles of known regulators of muscle morphogenesis on chromosome II, or to deficiencies that uncovered known regulators if lethal alleles were not publicly available. Screen lines that failed to complement adult viability were further analyzed for transheterozygous embryonic phenotypes. Only those lines that failed to complement embryonic phenotypes were classified as allelic to known regulators of muscle morphogenesis (see Table 2).

**Table 2.** Complementation groups.

| **Allele** | **Phenotype** |
| --- | --- |
| *arr*<sup>sj302</sup> | global morphogenesis, hypotrophic |
| *arr*<sup>sj359</sup> | global morphogenesis, hypotrophic |
| *barr*<sup>sj266</sup> | global morphogenesis |
| *barr*<sup>sj440</sup> | global morphogenesis, hypotrophic |
| *bsd*<sup>sj286</sup> | global morphogenesis |
| *bsd*<sup>sj348</sup> | VA2 missing |
| *bsd*<sup>sj362</sup> | global morphogenesis |
| *kon*<sup>sj11</sup> | DT1 LO1 VA3 missing |
| *kon*<sup>sj204</sup> | global morphogenesis |
| *kon*<sup>sj525</sup> | global morphogenesis |
| *ptc*<sup>sj287</sup> | global morphogenesis, hypertrophic |
| *ptc*<sup>sj461</sup> | global morphogenesis, hypertrophic |
| *salm*<sup>sj154</sup> | LO1 elongation and attachment |
| *salm*<sup>sj340</sup> | LO1 DT1 elongation and attachment |
| *ths*<sup>sj102</sup> | VA1 VA3 missing |
| *ths*<sup>sj237</sup> | VA2 elongation and attachment |
| *sj109* | global morphogenesis |
| *sj245* | global morphogenesis |
| *sj189* | VA3 elongation and attachment |
| *sj203* | VA2 elongation and branching |
|  | VA3 missing |
| *sj278* | VA3 missing |
| *sj264* | global morphogenesis |
| *sj455* | VA3 branching |
| *sj326* | hypotrophic muscles |
| *sj461* | hypotrophic muscles |

Stocks for complementation analysis included *bsd*^*1*^ (Yang et al., 2020a), *hoip*^*1*^ (Johnson et al., 2013), *kon*^*2986*^ (Johnson et al., 2013), *Df(2R)pyr36* (Kadam et al., 2009), *Df(2R)ths238* (Kadam et al., 2009), *arr*^*2*^, *barr*^*L305*^, *Df(2R)Egfr5, Df(2L)BSC354* [*eya*], *ptc*^*tuf-D*^, *Rca1*^*2*^, *robo*^*1*^, *S*^*IIN*^, *Sin3a*^*e64*^, *Sli*^*2*^, *smo*^*3*^, *Df(2R)BSC269* [*sns*], *tsh*^*04319*^, *tum*^*DH15*^, *and zip*^*1*^. Stocks were obtained from the Bloomington Stock Center unless otherwise referenced.

### Additional *Drosophila* stocks

Muscle morphology was visualized with *P{GMR40D04-GAL4}attP2* (*slou.Gal4*), *P{UAS-tdTom}*, and *P{UAS-eGFP}.* Myoblast fusion was assayed with *P{kirre*^*rP298*^*.lacZ}* (Nose et al., 1998). *Df(2R)Exel7098* was used for complementation analysis of putative *ptc* alleles. *Cyo, P{Gal4-Twi}, P{2X-UAS.eGFP}* was used to genotype embryos for screening, histology, and nucleic acid collection.

### Allele sequencing

For RNA deep sequencing (RNAseq), total RNA was collected from 7-12hr embryos per manufacturer’s specification (RNeasy kit, Qiagen), and submitted to Novogene (Sacramento, CA) for deep sequencing. cDNA libraries were prepped and sequenced using 150bp paired-end reads on the Illumina NovaSeq 6000 system. Two biological replicates were prepared and sequenced for 6 independent lines. Raw reads were quality and adapter trimmed using cutadapt (v2.4) as described (Martin, 2011). Trimmed reads were aligned using STAR (v2.5.4b) to the *Drosophila* reference genome BDGP6 with Ensemble Gene Annotation (v95) (Dobin et al., 2013). Variants for each line were called using aligned reads with the bcftools (v1.9) mpileup function followed by the bcftools call function (Li, 2011). Low quality variants were filtered using bcftools filter. Variants not found in both replicates from a line were discarded. Variants present in any complementary lines were discarded. Remaining variants were mapped to genes and prioritized using VEP (ensembl.org /Tools/VEP) (McLaren et al., 2016). RNAseq data and analysis are available at the Gene Expression Omnibus, accession number GSE164398. Causative variants were confirmed by complementation analysis with known alleles.

For Sanger sequencing, genomic DNA was collected from homozygous mutant 12-24hr embryos using the Quick Fly Genomic DNA prep method (BDGP). Overlapping PCR amplicons were generated across the coding sequences, and TOPO cloned into PCR2.1 (Thermofisher). M13R and T7 primers were used to sequence inserts from both the 5’ and 3’ ends. Point mutations were validated by sequencing multiple PCR products.

### Immunohistochemistry and imaging

α-βgal (1:100, Promega, Z3781) was used to visualize rP298.lacZ. Embryo antibody staining was performed as described (Johnson et al., 2013); primary antibodies were detected with HRP-conjugated secondary antibodies and the TSA system (Molecular Probes). Embryos from the primary screen were live imaged on an Axio Observer 5 compound fluorescent microscope; follow-up imaging was used a Zeiss LSM800 confocal microscope. Imaging parameters were maintained between genotypes where possible.

### Data and reagent availability

All data necessary for confirming the conclusions of this article are represented fully within the article and its tables and figures. All fly stocks are available upon request.

## Results and Discussion

An invariant number of mesodermal founder myoblasts and ectodermal tendon precursors (tenocytes) are specified in each abdominal segment (A2-A8) of the *Drosophila* embryo (de Joussineau et al., 2012; Frommer et al., 1996). By the end of myogenesis, the two deterministic cell populations locate each other with a precise spatiotemporal stoichiometry, ostensibly through the process of myotube guidance (Fig. 1A,B)(Yang et al., 2020b). To better understand myotube guidance at the genetic level, we screened for novel regulators of muscle morphogenesis using an EMS-based forward genetic approach (Fig. 1C).

Using *slou>tdTomato*, which is expressed in 5 founder myoblasts, we screened 564 embryonic lethal alleles on the second chromosome for myogenic phenotypes. Our screen produced 58 alleles that grouped into 7 classes including morphogenesis defects in all 5 muscles (Fig. 2A), morphogenesis defects in a subset of muscles (Fig. 2B-D), hypotrophic and hypertrophic muscles (Table 1), and missing muscles (Table 1). Complementation analysis among the 58 alleles identified 11 complementation groups with multiple alleles, and 33 complementation groups with just a single allele (Table 2). In total our screen identified 44 loci that regulate muscle morphogenesis, and alleles in 32 loci appeared to cause specific phenotypes. Alleles of the remaining 12 genes appeared to cause severe phenotypes that disrupted anterior-posterior patterning, germ band retraction, or dorsal closure.

**Figure 2.**
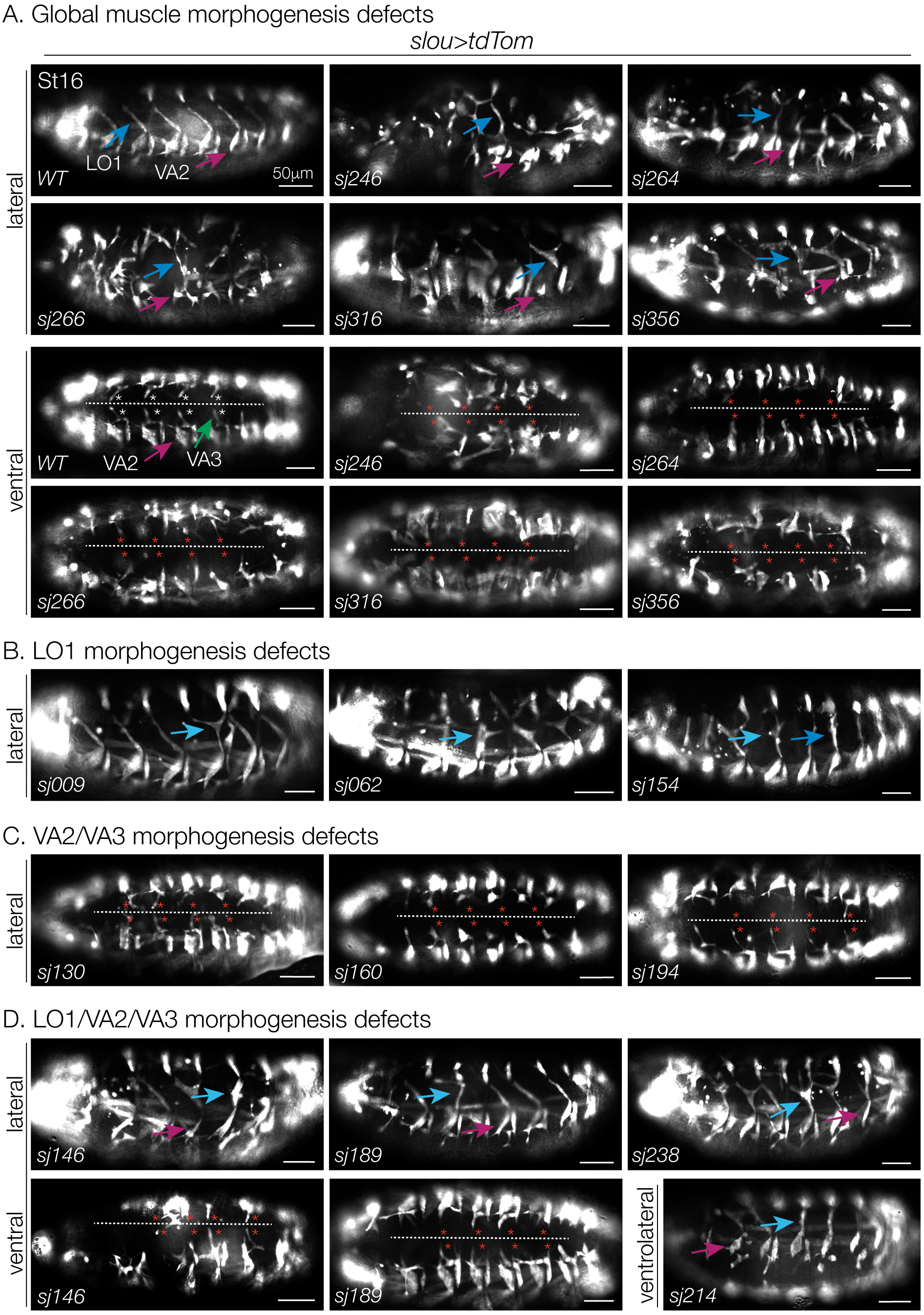
Phenotypic classes. Live Stage 16 embryos that expressed *slou>tdTomato*. In lateral views, LO1 (blue arrows) and VA2 (magenta arrows) muscles are indicated. In ventral views, the ventral midline (dotted line) and the expected positions of VA3 attachment sites (asterisks) are highlighted. (A) Global morphogenesis defects. The shape or position of all 5 muscles (LO1, DT1, VT1, VA2, and VA3) were affected. (B) LO1 morphogenesis defects. Oblique morphology often transformed to longitudinal morphology. (C) VA2/VA3 morphogenesis defects. VA3 muscles frequently failed to attach near the ventral midline (red asterisks). (D) LO1/VA2/VA3 morphogenesis defects. Muscle phenotypes similar to (B,C). Notice VA3 crosses the ventral midline in *sj146*.

### 26 novel regulators of muscle morphogenesis

We have characterized 4 genes on the second chromosome that regulate myotube guidance including *hoi polloi* (*hoip*)(Johnson et al., 2013), *pyramus* (*pyr*) and *thisbe* (*ths*) (Yang et al., 2020b), and *back seat driver* (*bsd*) (Yang et al., 2020a). *kon tiki* (*kon*) (Schnorrer et al., 2007), *slit*, *robo* (Kramer et al., 2001), and *tumbleweed* (*tum*)(Guerin and Kramer, 2009; Yang et al., 2020a) are also located on chromosome II and direct myotube guidance. In addition, *arrow* (*arr*), *barren* (*barr*), *Epidermal growth factor receptor* (*Egfr), eyes absent* (*eya*), *patched (ptc), Regulator of cyclin A1 (Rca1), Star (S), Sin3a, smoothened (smo)*, *teashirt* (*tsh*), and *zipper* (*zip*) were identified in a previous screen for regulators of muscle morphogenesis on the second chromosome (Chen et al., 2008). Thus a total of 19 genes involved in muscle morphogenesis have been mapped to the second chromosome.

Complementation analysis between the 58 alleles recovered in our screen and alleles of the 19 known regulators of muscle morphogenesis showed that our screen identified 13 new alleles in 6 known morphogenetic loci (see Materials and Methods for allele nomenclature). Two alleles mapped to *bsd*, which encodes a ser/thr kinase (Fig. 3A), and by Sanger sequencing we found *bsd*^*s286*^ is a missense mutation in the Bsd ATP binding pocket and *bsd*^*sj362*^ is a missense mutation in the Bsd kinase domain. Our previous study suggested Bsd kinase activity is required for accurate myotube guidance (Yang et al., 2020a), and the identification of new alleles that affect the Bsd kinase domain strengthens this hypothesis. We also identified new alleles of *ths*, *kon*, *arr*, *barr*, and *tum* (Figs. 3B,C Fig. 4B,C Table 2). Alleles in the remaining 26 loci fully complemented all known regulators of muscle morphogenesis, suggesting the function of at least 26 genes remains to be characterized in the context of myogenesis (Tables 1,2).

**Figure 3.**
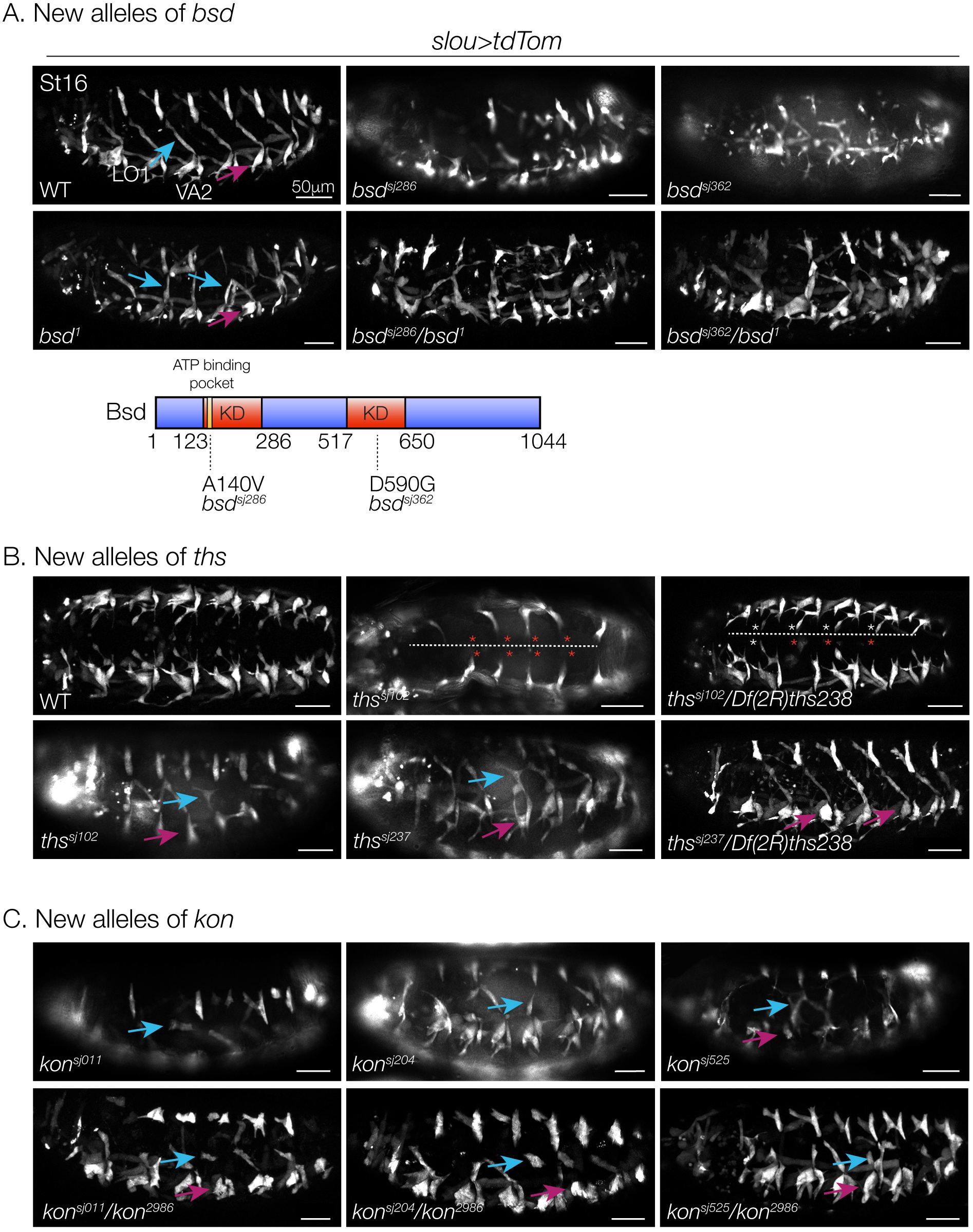
New alleles in known regulators of myotube guidance. Live Stage 16 embryos that expressed *slou>tdTomato* labeled as in Fig. 2. (A) *bsd*^*sj286*^ and *bsd*^*sj362*^ failed to complement *bsd*^*1*^. A schematic of the Bsd protein is shown; *bsd*^*sj286*^ is a A140V missense mutation in the ATP binding domain and *bsd*^*sj362*^ is a D590G missense mutation in the kinase domain (KD). (B) *ths*^*sj102*^ and *ths*^*sj237*^ failed to complement *Df(2R)ths238*. VA3 muscles that failed to attach near the ventral midline are indicated (red asterisks). (C) *kon*^*sj11*^, *kon*^*sj204*^, and *kon*^*sj525*^ failed to complement *kon*^*2986*^. LO1 muscles (blue arrows) were often rounded, indicative of myotube elongation defects.

**Figure 4.**
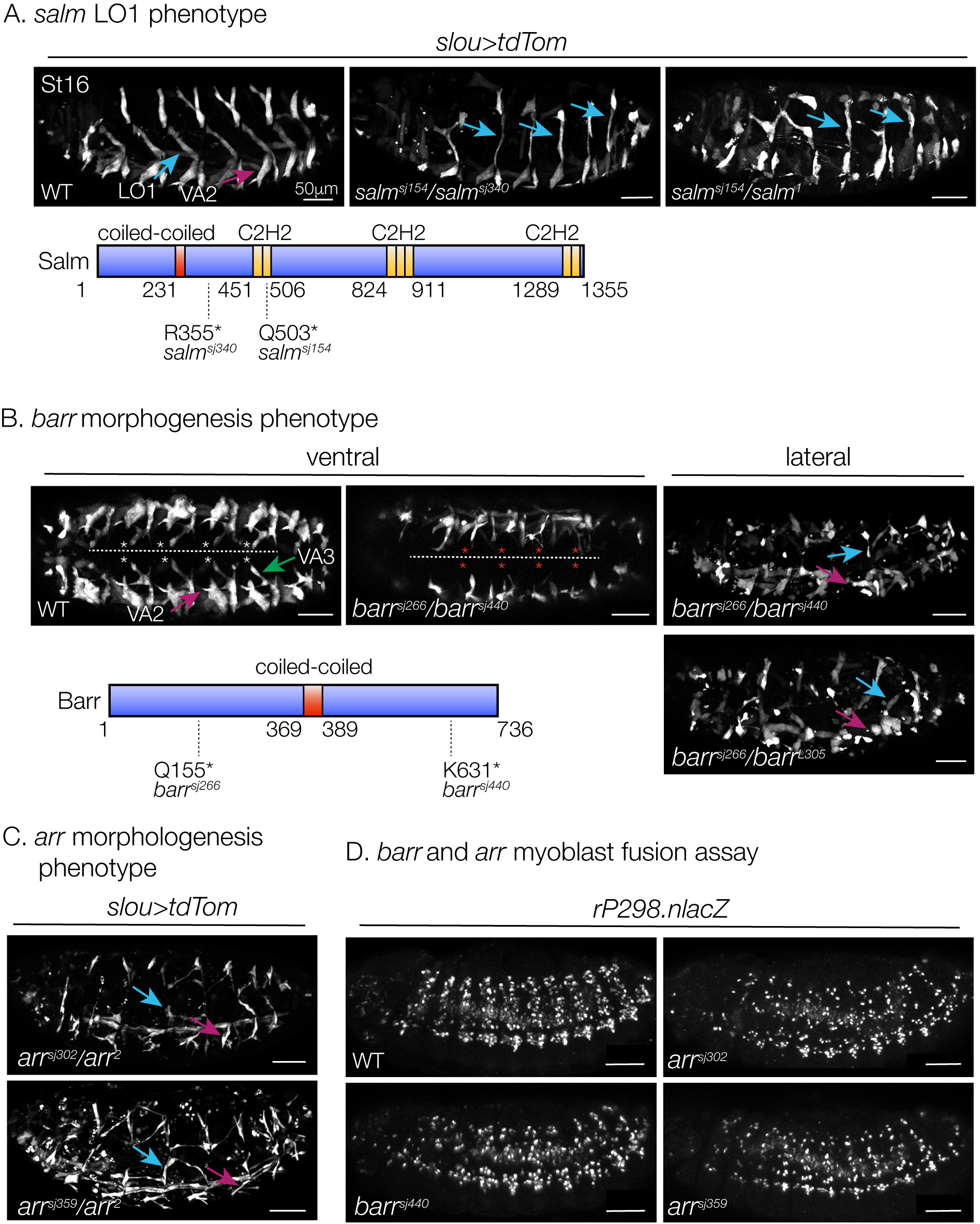
Salm, Barr, and Arr direct muscle morphogenesis. (A-C) Live Stage 16 embryos that expressed *slou>tdTomato* labeled as in Fig. 2. (A) *salm*^*sj154*^/*salm*^*sj340*^ embryos phenocopied the myogenic defects of *salm*^*sj154*^/*salm*^*1*^ embryos. Note the longitudinal morphology of LO1 muscles (blue arrows). A schematic of the Salm protein is shown; *salm*^*sj154*^ is a Q503* nonsense mutation and *salm*^*sj340*^ is a R355* nonsense mutation. Yellow boxes represent the 7 C2H2-type zinc finger domains (C2H2). (B) VA3 muscles showed attachment site defects in *barr*^*sj266*^*/barr*^*sj440*^ embryos (ventral view, red asterisks). *barr*^*sj266*^*/barr*^*sj440*^ embryos phenocopied *barr*^*sj266*^*/barr*^*L305*^ embryos (lateral view). The Q155* and K631* nonsense mutations are shown on the schematic of the Barr protein. (C) *arr*^*sj302*^ and *arr*^*sj359*^ embryos showed LO1 muscle (blue arrows) and VA2 muscle (magenta arrows) morphogenesis phenotypes. (D) Myoblast fusion assay. Stage 12 embryos labeled for rP298.nlacZ, which identifies myogenic nuclei. *barr*^*sj440*^, *arr*^*sj302*^, and *arr*^*sj359*^ embryos showed fewer lacZ+ nuclei after the first round of myoblast fusion.

### Mapping unique phenotypes

Morphogenetic studies require a toolkit that is often distinct from studies of cell fate specification since morphogenesis phenotypes are rarely quantified by the simple presence or absence of a specific transcript, and multiple morphogenetic events can occur simultaneously. From this perspective, we chose to follow-up on (1) alleles that produced an unusually specific phenotype and (2) alleles of a single locus that produced distinct phenotypes. In addition to the 26 uncharacterized loci we identified, the mechanisms by which *arr, barr, ptc, Rca1, smo*, *tsh*, and *zip* regulate myogenesis are unknown. Among these 33 loci, 3 sets of allelic mutations produced unique phenotypes with respect to myogenesis.

*sj154* and *sj340* mutant embryos showed a specific phenotype in which the Longitudinal Oblique 1 (LO1) muscle showed abnormal morphology, while the remaining muscles looked normal (Fig. 2B). *barr*^*sj266*^ and *barr*^*sj440*^ embryos showed global muscle morphogenesis defects, but *barr*^*sj440*^ embryos had thin, hypotrophic muscles while *barr*^*sj266*^ embryos had muscles of normal size (Fig. 4B). *sj287* and *sj461* also caused global muscle morphogenesis defects, but the mutant muscles were hypertrophic (Fig. 5A). We identified 6 other alleles that caused hypotrophic muscles (Table 1), but *sj287* and *sj461* were the only alleles that caused a hypertrophic muscle phenotype.

**Figure 5.**
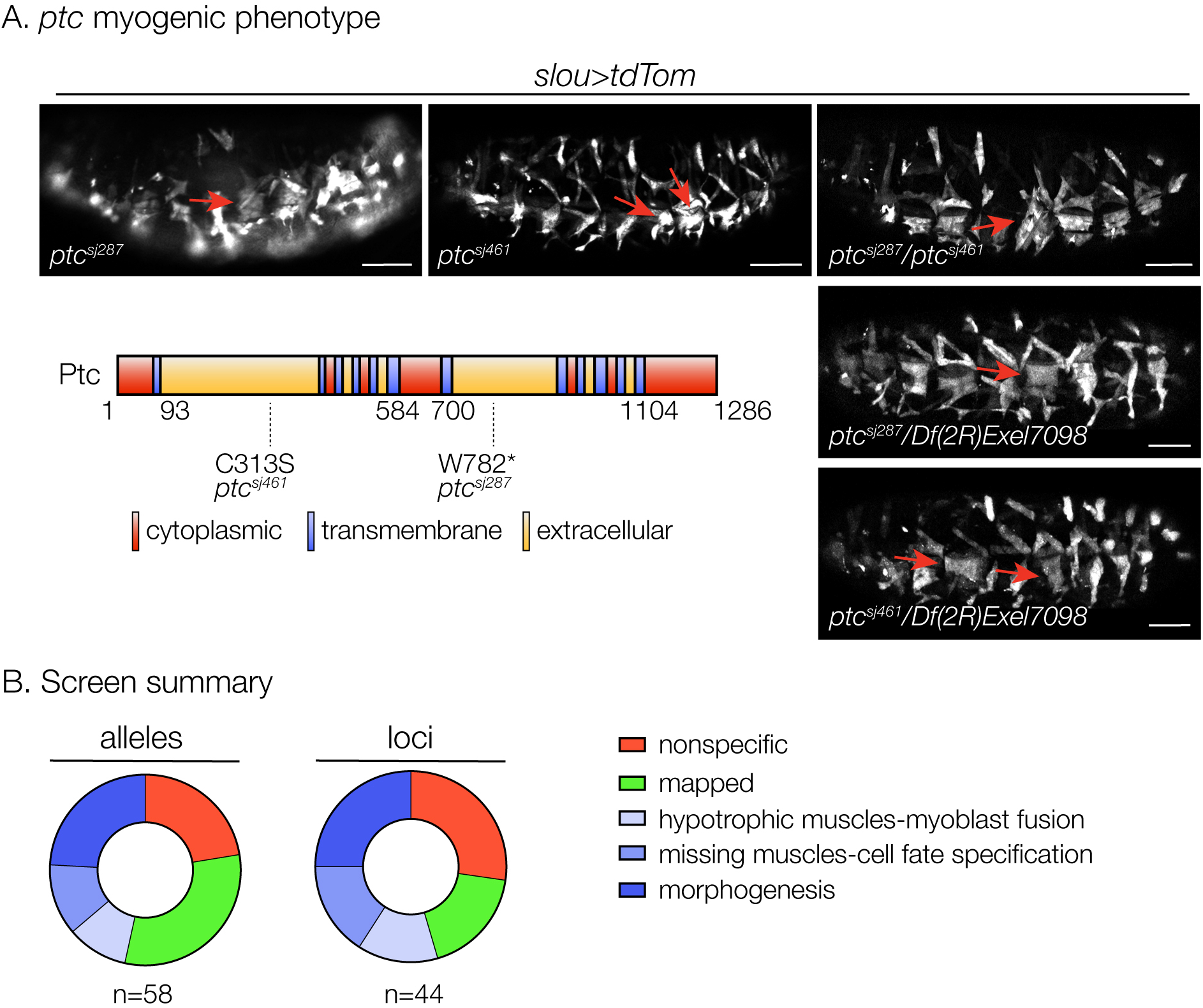
Hh signaling regulates muscle size. (A) Live Stage 16 embryos that expressed *slou>tdTomato*. Muscles in *ptc*^*sj287*^ and *ptc*^*sj461*^ embryos were hypertrophic, particularly VA2 (red arrows). *ptc*^*sj287*^/*ptc*^*sj461*^ transheterozygous embryos phenocopied *ptc*^*sj287*^/*Df(2R)Exel7098 and ptc*^*sj461*^/*Df(2R)Exel7098* embryos. A schematic of the Ptc protein is shown with 12 transmembrane domains; *ptc*^*sj461*^ is a C313S missense mutation in an extracellular domain and *ptc*^*sj287*^ is a nonsense mutation. (B) Screen results. 58 alleles in 44 loci were identified that affect muscle morphogenesis. Greater than half of the alleles were mapped or found to be nonspecific (e.g. anterior-posterior patterning, germ band retraction, dorsal closure). Unmapped alleles with specific phenotypes are shaded blue. Alleles producing hypotrophic muscles were shown to have defects in myoblast fusion; alleles that cause missing muscles are presumed to affect cell fate specification. 14 alleles (in 11 loci) that disrupt muscle morphogenesis (dark blue) remain to be mapped.

We mapped *barr*^*sj266*^, *barr*^*sj440*^, *sj154, sj340, sj287*, and *sj461* by RNA deep sequencing (RNAseq), and confirmed *barr*^*sj266*^ and *barr*^*sj440*^ are new alleles of *barr* (Fig. 4B). *sj154* and *sj340* mapped to nonsense mutations in *salm*, and *sj287* and *sj461* mapped to a missense and a nonsense mutation in *ptc* (Figs. 4A, 5A). To understand if the sequenced mutations were causative of myogenic phenotypes, we performed complementation tests with known alleles, and in all 6 cases we found transheterozygous embryos phenocopied the myogenic defects of the sequenced homozygotes (Figs 4A,B and 5A).

### Salm directs embryonic muscle morphogenesis

LO1 muscles in *salm*^*sj154*^/*salm*^*sj340*^ embryos showed longitudinal instead of oblique morphology (Fig. 4A). *salm* encodes a zing finger transcription factor that controls fiber type identity during adult myogenesis (Bryantsev et al., 2012; Schönbauer et al., 2011), but a role for *salm* in embryonic muscle morphogenesis has not been characterized. Our data suggest Salm could be a transcriptional regulator of myotube guidance.

### Epigenetic regulation of myoblast fusion and myotube guidance

All 5 *slou.Gal4* positive muscles in *barr*^*sj266*^/*barr*^*sj440*^ embryos were misshapen (Fig. 4B). Although *barr* was identified in a previous muscle screen (Chen et al., 2008), the role of Barr during myogenesis is unknown. Barr is a chromatin binding protein and is part of the condensin complex that directs chromosome condensation, which is a landmark of early mitosis (Herzog et al., 2013). Since founder cells and fusion competent myoblasts exit the cell cycle prior to myoblast fusion and myotube guidance, it is possible that Barr regulates chromatin for a mitotic-independent function during muscle morphogenesis.

We were initially interested in *barr*^*sj266*^ and *barr*^*sj440*^ because *barr*^*sj440*^ muscles were hypotrophic while *barr*^*sj266*^ muscles were normal in size. Hypotrophic muscles are generally indicative of myoblast fusion defects, so we assayed myoblast fusion in all of the hypotrophic lines we recovered in the screen (*sj15, sj302, sj314, sj326, arr*^*sj302*^, *arr*^*sj359*^, and *barr*^*sj440*^). The first round of myoblast fusion was visibly disrupted in all lines except *sj15* (Fig. 4D and data not shown), suggesting hypotrophic muscles were indeed the result of myoblast fusion defects. These data argue that Barr regulates myoblast fusion and myotube guidance, and highlight the exciting possibility that the *barr*^*sj266*^ and *barr*^*sj440*^ alleles can be used to genetically separate the role of Barr in myoblast fusion from its role in myotube guidance. Such studies will likely reveal a novel epigenetic role for Barr outside of chromosome condensation.

The transmembrane receptor Arr, which modulates Wingless (Wg) signaling, is also required for myoblast fusion and presumably for myotube guidance (Fig. 4C,D). We had previously shown that Wg signaling regulates myoblast specification (Duan et al., 2001), however embryos homozygous for *arr*^*sj302*^ and *arr*^*sj359*^ appeared to show correct founder cell fate specification but defective myotube morphogenesis (Fig. 4C). The *arr* alleles may provide future insights into the role of Wg signaling in regulating muscle morphogenesis downstream of cell fate specification.

### Hh signaling dictates muscle size

*ptc*^*sj287*^ and *ptc*^*sj461*^ embryos showed hypertrophic muscles, but both lines complemented the lethal allele *ptc*^*tuf-D*^ in our initial complementation analysis. Since our RNAseq analysis of the hypertrophic lines identified deleterious mutations in the *ptc* coding region, we assayed muscle morphogenesis in embryos transheterozygous for *Df(2R)Exel7098* which uncovers *ptc*. *ptc*^*sj287*^*/Df(2R)Exel7098* and *ptc*^*sj461*^*/Df(2R)Exel7098* embryos had hypertrophic muscles, confirming the *ptc* coding region mutations are causative of the muscle phenotype.

The mechanisms that promote muscle hypertrophy in neonatal, adolescent, and adult vertebrates have been studied in great detail (Sartori et al., 2021). What is less appreciated is that muscle size must also be regulated in the developing embryo, and that unknown mechanisms are in place to limit muscle overgrowth. To our knowledge, the *ptc*^*sj287*^ and *ptc*^*sj461*^ embryos are the first examples of hypertrophic embryonic muscles, and may provide a starting point for understanding how excessive muscle hypertrophy is limited during development. *ptc*^*sj287*^ and *ptc*^*sj461*^ myotubes also showed phenotypes consistent with myotube guidance defects, suggesting Hh signals provide guidance cues as well as growth regulatory information during *Drosophila* myogenesis.

### Toward a molecular understanding of muscle topography

Our working hypothesis is that muscle topography is established through the actions of short-range secreted signals that provide navigational cues to growing myotubes, and tenocyte expressed cell recognition molecules that direct myotubes to select the correct muscle attachment site. The results from this screen warrant further investigation into Wg and Hh signals as navigational cues during myotube guidance, and provide new clues into the gene regulatory events, controlled by Salm and Barr, that coordinate cellular guidance with attachment site selection. We successfully used RNAseq to map 6 new alleles without the laborious recombination mapping usually needed to map chemically induced point mutations. 13 new alleles were also mapped to known regulators of muscle morphogenesis. However, the mapped alleles represent just a fraction of the mutations with specific phenotypes that we recovered from this screen (Fig. 5B), arguing many additional loci remain to be identified that regulate muscle morphogenesis.

## Acknowledgements

We thank all members of the Johnson and Skeath labs for their encouragement and support. ANJ was supported by NIH R01AR070299 (NIAMS), and JBS was supported by NIH R01NS036570 (NINDS).

